# Elasmobranch genome sequencing reveals evolutionary trends of vertebrate karyotype organization

**DOI:** 10.1101/2022.10.17.512540

**Authors:** Kazuaki Yamaguchi, Yoshinobu Uno, Mitsutaka Kadota, Osamu Nishimura, Ryo Nozu, Kiyomi Murakumo, Rui Matsumoto, Keiichi Sato, Shigehiro Kuraku

## Abstract

Genomic studies of vertebrate chromosome evolution have long been hindered by the scarcity of chromosome-scale DNA sequences of some key taxa. One of those limiting taxa has been the elasmobranchs (sharks and rays), which harbor species often with numerous chromosomes and enlarged genomes. Here, we report the chromosome-scale genome assembly for the zebra shark *Stegostoma tigrinum*, an endangered species that has the smallest genome sequenced to date among sharks (3.71 Gb), as well as for the whale shark *Rhincodon typus*. Our analysis employing a male–female comparison identified an X chromosome, the first genomically characterized shark sex chromosome. The X chromosome harbors a Hox C cluster whose intact linkage has not been shown for an elasmobranch fish. The sequenced shark genomes exhibit a gradualism of chromosome length with remarkable length-dependent characteristics—shorter chromosomes tend to have higher GC content, gene density, synonymous substitution rate, and simple tandem repeat content as well as smaller gene length, which resemble the edges of longer chromosomes. This pattern of intragenomic heterogeneity, previously recognized as peculiar to species with so-called microchromosomes, occurs in more vertebrates including elasmobranchs. We challenge the traditional binary classification of karyotypes as with and without microchromosomes, as even without microchromosomes, shorter chromosomes tend to have higher contents of GC and simple tandem repeats and harbor shorter and more rapid-evolving genes. Such characteristics also appear on the edges of longer chromosomes. Our investigation of elasmobranch karyotypes underpins their unique characteristics and provides clues for understanding how vertebrate karyotypes accommodate intragenomic heterogeneity to realize a complex readout.

## Introduction

Genomes accommodate coexisting regions with differential characteristics, and these characteristics are manifested not only in DNA sequences (e.g., GC-content) but also in intra-genomic heterogeneity of non-sequence features such as chromatin openness, replication timing, and recombination frequency. These features are thought to be associated with how karyotypes of individual species are organized.

For example, in the chicken genome, early replicating regions tend to be found in small chromosomes, known as ‘microchromosomes’ (see below). It is still unknown how such intragenomic heterogeneity is accommodated by variable karyotypes as well as how it arose during evolution. To reconstruct the ancestor of all extant vertebrates and the evolutionary process thereafter, information from the evolutionary lineages that branched off in the early phase of vertebrate evolution is instrumental. Among those lineages, whole genome sequence information for cartilaginous fishes has been scarce. The importance of studying cartilaginous fishes is doubled when we consider their genomic trends. No additional whole genome duplication has been reported for cartilaginous fishes, while drastic lineage-specific genomic changes are being reported for the other extant non-osteichthyan taxon, cyclostomes (Nakatani et al. 2021).

Cartilaginous fishes (chondrichthyans) are divided into two groups, Holocephala (chimaera and ratfish) and Elasmobranchii (sharks and rays). Even long after whole genome sequences of a holocephalan species, *Callorhinuchus milii*, were made available (Venkatesh et al. 2014), those sequences have not been validated with any karyotyping report. This limitation stems mainly from the technical difficulty in reproducibly preparing chromosome spreads from a stable supply of metaphase cells. Only recently has our repeated sampling of fresh shark tissues (blood or embryos) (e.g., at aquariums) enabled karyotyping using culture cells for four orectolobiform shark species in Elasmobranchii (Uno et al. 2020). This has paved the way for rigid evaluation of whole genome sequences. Biological studies on cartilaginous fishes have been hindered by low accessibility to fresh material. Moreover, especially in studying elasmobranchs, genome analysis can encounter inherent difficulty incurred by their large genome sizes (reviewed in Kuraku 2021). These factors have prevented previous efforts on cartilaginous fishes from obtaining a suite of genome sequences supported by karyotype and genome size estimate as well as transcriptome sequencing (Rhie et al. 2021; Read et al. 2017; Tan et al. 2021; Weber et al. 2020; Marra et al. 2019; Zhang et al. 2020).

Chromosome-level analysis is broadening our scope of comparative genomics (Rhie et al. 2021; Deakin et al. 2019). The difficulty of elasmobranch genome sequencing is manifested in the retrieval of the Hox C cluster, an array of homeobox-containing genes (reviewed in Kuraku 2021). While shark Hox A, B, and D gene clusters have been reliably assembled with few gaps (Hara et al. 2018; Mulley et al. 2009), assembling the Hox C cluster, which was initially reported as missing from elasmobranch genomes (King et al. 2011), suffered from high GC-content and frequent repetitive elements, resulting in fragmentary sequences (Hara et al. 2018; reviewed in Kuraku 2021). This difficulty is expected to be overcome by the application of a rising approach to scaffolding the genomic fragments up to the chromosome scale using chromatin contact data (Yamaguchi et al. 2021; Dudchenko et al. 2017) as well as long-read sequencing.

Another expectation from chromosome-level genome analysis is the identification of sex chromosomes, although sequencing and assembling sex chromosomes often suffer from difficulties caused by high repetitiveness or uneven sequence depth (Rhie et al. 2021; Ma et al. 2021). While sex determination mechanisms have been revealed by an increasing number of studies on osteichthyan vertebrates (Graves 2016; Pennell et al. 2018), no report is available for vertebrate species outside osteichthyans, namely cyclostomes and chondrichthyans. The quest for sex determination mechanisms can be initiated by the identification of sex chromosomes as already improvised in several vertebrate species (Franchini et al. 2018).

Some vertebrate karyotypes, including most bird karyotypes, consist of small-sized chromosomes, or microchromosomes (Figure 1A) that were initially recognized as shorter than 1 μm in cytogenetic observations (Ohno et al. 1969; Ohno 1970). Microchromosomes are known to have higher GC-content, higher gene density, and different chromatin states compared with the remaining macrochromosomes (International Chicken Genome Sequencing Consortium (ICGSC) 2004; Burt 2002; Waters et al. 2021). Some genome sequence-based studies regard chromosomes smaller than 20 Mb as microchromosomes and investigated their possible common origin (Nakatani et al. 2021; ICGSC 2004), but they have not been unambiguously defined on a cross-species basis. Accumulating information from synteny-based analysis suggests that the last common jawed vertebrate ancestor already possessed microchromosomes (Braasch et al. 2016; Waters et al. 2021; Nakatani et al. 2007, 2021). However, this hypothesis needs to be examined by incorporating more diverse vertebrate taxa into the comparison, on a solid basis of experimentally validated karyotypic configurations of individual species.

**Figure 1.**
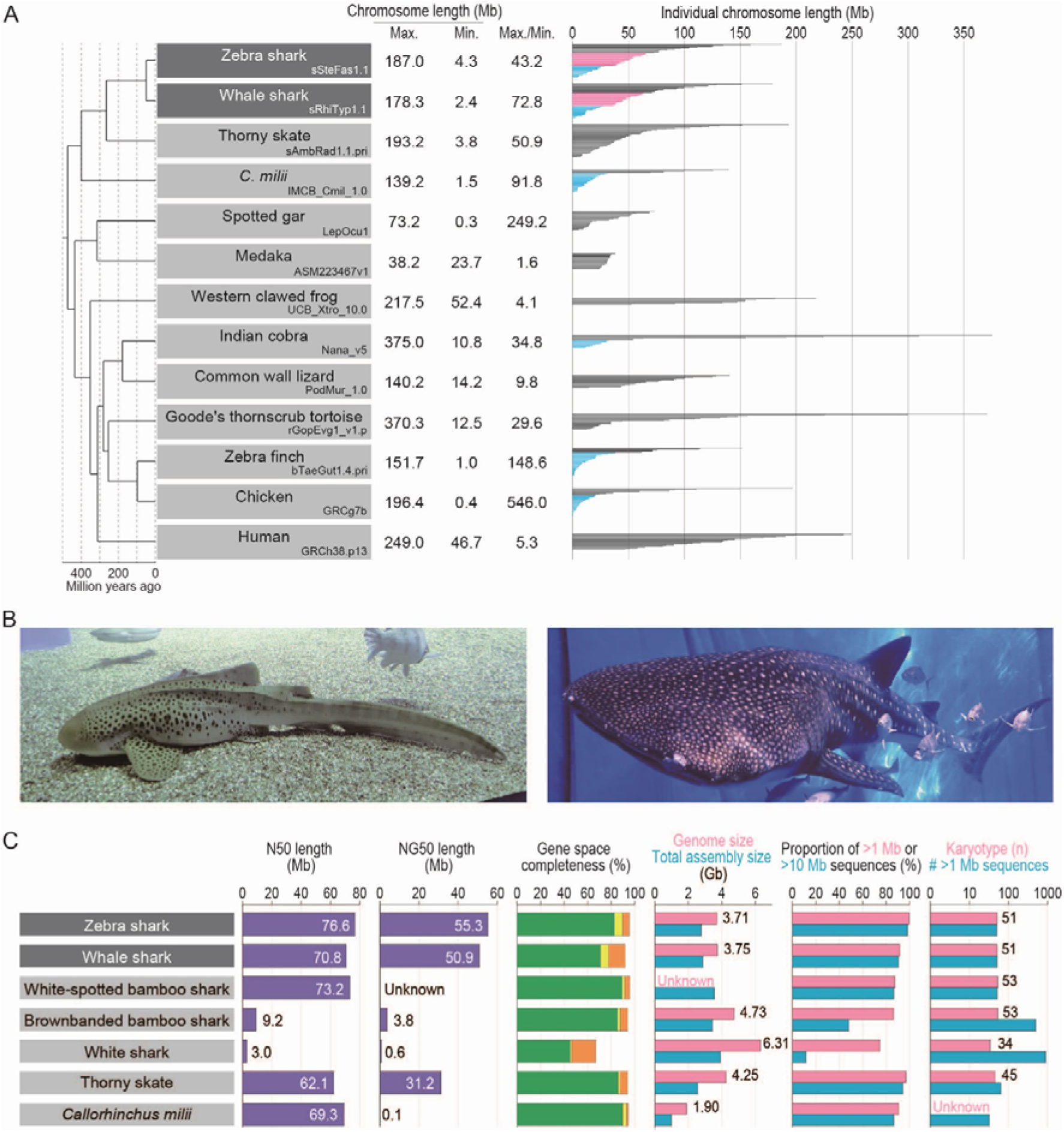
Shark species studied and comparative statistics of their genome assemblies. (*A*) The karyotypes of diverse vertebrate species are depicted with the length of individual chromosomal DNA sequences. The maximum and minimum chromosome lengths are based on the records in NCBI Genomes. Microchromosomes for osteichthyans are shown in light blue according to individual original reports (Nakatani et al. 2021; Suryamohan et al. 2020; Knief and Forstmeier 2016; ICGSC 2004), while the eMID and eMIC of the two shark species whose genomes have been sequenced in the present study (see Results) are shown in orange and light blue, respectively. (*B*) Zebra shark (left) and whale shark (right). (*C*) Statistics of the genome assemblies. The identifiers of the genome assemblies, as well as statistical comparison of more metrics, are included in Supplementary Table 1. Gene space completeness shows the proportions of selected one-to-one protein-coding orthologs with ‘complete’ (green), ‘duplicated’ (yellow), and ‘fragmented’ (orange) coverages retrieved in the sequences by the BUSCO pipeline (see Methods). The details of the genome sizes and karyotypes included are based on existing literature (Uno et al. 2020; Schwartz and Maddock 1986, 2002; Hardie and Hebert 2004).

In this study, we focused on the zebra shark *Stegostoma tigrinum* (or leopard shark; Figure 1B) and report its whole-genome sequences for the first time. This species has the smallest genome size (3.71 Gb) among the elasmobranch species whose genomes have been sequenced to date. Each of the resultant chromosome-scale genome assemblies of the zebra shark, as well as the whale shark (Figure 1B), have been constructed using samples from a single individual (which was not achieved in earlier efforts: Tan et al. 2021; Read et al. 2017; Weber et al. 2020) and controlled by referring to their karyotypes. Using the obtained sequences, we performed comparative investigations to characterize the diversity of chromosomal organization.

## Results

### The smallest shark genome sequenced to date

We focused on the zebra shark (or leopard shark) *Stegostoma tigrinum* (formerly, *S. fasciatum*) with the haploid nuclear DNA content of 3.79 pg (3.71 Gb; Kadota et al. in prep) and the karyotype of 2n = 102 (Uno et al. 2020). Using genomic DNA extracted from blood cells of a female adult, the whole genome was sequenced and assembled with short reads. The resultant sequences were further scaffolded using Hi-C data obtained from blood cell nuclei of the same individual (see Methods). The number of output scaffold sequences longer than 1 Mbp (thereafter tentatively designated ‘chromosomes’) was 50, which closely approximates its chromosome number revealed by the cytogenetic observation and karyotypic organization previously characterized by us using primary cultured cells (Figure 1C). The retrieval of the chromosomal scale was also guaranteed by the N50 scaffold length of 76.6 Mb (Figure 1C).

We also performed *de novo* whole genome sequencing of an adult male whale shark, *Rhincodon typus*, using Linked-Read data and Hi-C scaffolding (see Methods). The number of sequences greater than 1 Mbp matched the number of chromosomes in the karyotype (n = 51) (Figure 1C), and the N50 scaffold length of the resultant assembly reached 70.8 Mb, significantly exceeding that of the assemblies previously published for this species (Figure 1C). To date, our product is the only chromosome-scale genome assembly for this species that was built consistently from a single individual.

While we recognize a significant gap between the estimated and retrieved total sequence lengths, genome assemblies for the zebra shark and whale shark exhibited high completeness of protein-coding gene space of more than 90% (Figure 1C). Prediction of protein-coding genes on the zebra shark and whale shark genomes, performed by incorporating homolog sequences and transcriptome data, resulted in 33,222 and 35,334 genes, respectively (Supplementary Tables 1 and 2), which allowed downstream molecular biological analysis.

### Karyotypic trends in sharks

The obtained zebra shark genome assembly consisted of chromosome sequences of highly variable length with a gradual slope, in accordance with our previous cytogenetic observation (Uno et al. 2020), spanning from 187.0 Mb down to 4.3 Mb (Figure 2A). This pattern does not resemble the length variation in the *Callorhinchus milii* or the chicken (Figure 2A). The chicken especially exhibits a steep slope, marked by a number of chromosomes shorter than 20 Mb (microchromosomes) (Figures 1A and 2A). Gradualism in the chromosome length distribution is also observed in the whale shark (Supplementary Figure 1), white-spotted bamboo shark (Zhang et al. 2020), and thorny skate (Rhie et al. 2021), and is assumed to be typical karyotypic organization of elasmobranchs (Supplementary Figure 2). For simplicity, we tentatively designated (1) zebra shark chromosomes 1 to 14, longer than 70 Mb; (2) chromosomes 15 to 33, between 30 and 70 Mb; and (3) chromosomes 34 to 50, shorter than 30 Mb as <u>e</u>lasmobranch <u>mac</u>ro-, <u>mid</u>dle-sized-, and <u>mic</u>ro-chromosomes (abbreviated into eMAC, eMID, and eMIC, respectively) that are differentially colored in Figures 1A and 2B. This categorization also applies directly to the whale shark chromosomes (Figure 1A; Supplementary Figure 2).

**Figure 2.**
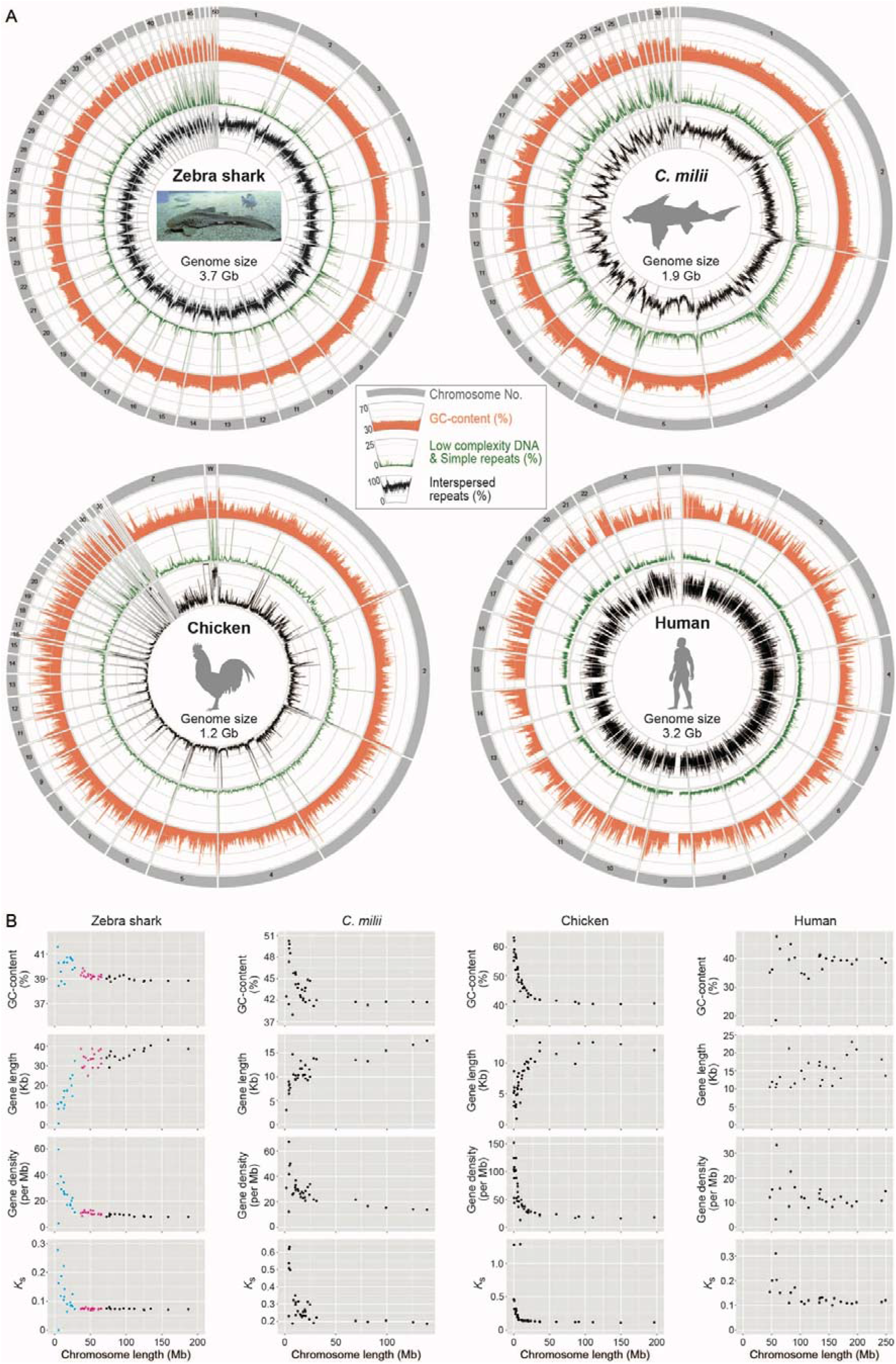
Chromosomal sequence compositions of the zebra shark and selected vertebrate species. (*A*) Sequence characterization of different chromosomal segments. The orange areas show GC-content (30%–70%), while the green and black lines show content of simple tandem repeats (0%–25%) and interspersed repeats (0%–100%), respectively, in 100-kb-long non-overlapping windows. (*B*) Two-dimensional plots of GC-content and median values of gene length, gene density, and the median of synonymous substitutions per synonymous site (*K*_s_) for protein-coding genes on individual chromosomes. Computation of *K*_s_ was performed for each of the four species in order by involving a pair species, namely, whale shark *Rhincodon typus*; small-eyed rabbitfish *Hydrolagus affinis*; helmeted guineafowl *Numida meleagris*; and common marmoset *Callithrix jacchus*. Zebra shark chromosomes are shown as three groups, eMAC, eMID, and eMIC (see text for details), in black, magenta, and cyan, respectively.

The zebra shark genome exhibits consistent intra-chromosomal GC-content compared with other species, and its chromosome ends in general, as well as the shorter chromosomes, tend to have relatively higher GC-content (Figure 2A). Zebra shark DNA sequences show a uniformly high frequency of interspersed repeats throughout the genome, while its shorter chromosomes tend to have higher simple tandem repeat content (Figure 2A). These characteristics are also observed in the whale shark genome (Supplementary Figure 1), but the co-occurrence of these two features was not explicitly observed in the other species compared (Figure 2A; Supplementary Figure 1).

### Vertebrate-wide comparisons including elasmobranchs

To analyze how the karyotypes of these shark species were derived, chromosomal nucleotide sequences were compared between species pairs with variable divergence times (Figure 3A). The comparison between the zebra shark and the whale shark exhibited a high similarity in chromosomal organization with few intrachromosomal breaks (panel 1 in Figure 3A). The high similarity of this orectolobiform shark species pair suggests high conservation of genomic sequences from around or earlier than 50 million years ago (see Discussion), compared with osteichthyan species pairs in a similar divergence time range-the human-marmoset pair diverged about 43 million years ago (panel 3 in Figure 3A). The relatively high conservation of chondrichthyan chromosome organization is also supported by the comparisons between more distantly related species pairs. The similarity of the zebra shark genome sequences to the thorny skate (panel 4) and the *C. milii* (panel 6) exceeded those for the species pairs with about 300- and 400-million-year divergences, respectively (panel 5 and panels 7 in Figure 3A).

**Figure 3.**
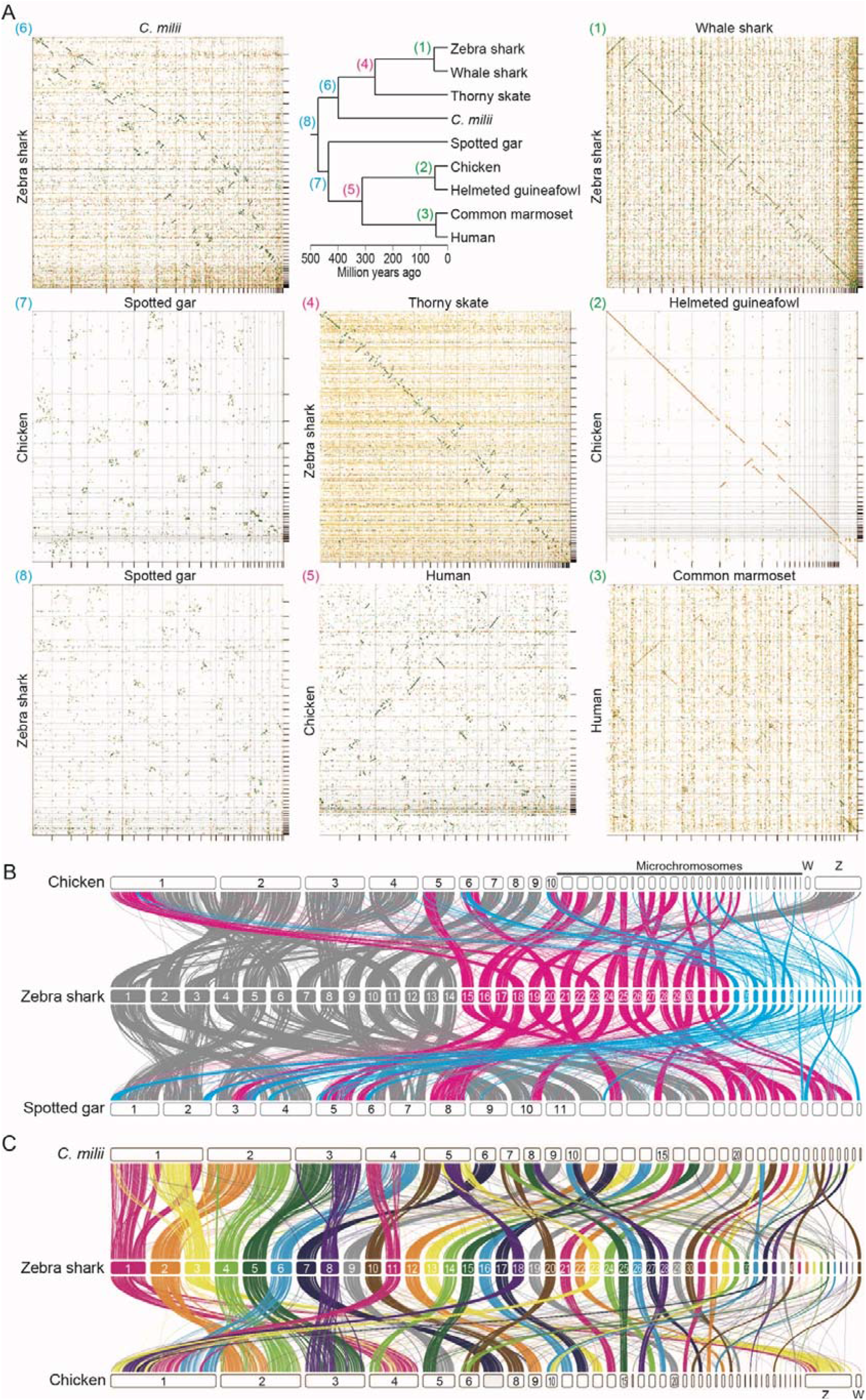
Cross-species investigation of chromosomal homology. (*A*) Dot matrices showing genome sequence similarities for selected pairs of vertebrate species with variable divergence times. Sequences of high similarity are shown with diagonal lines by the program D-GENIES (Cabanettes and Klopp 2018). The numbers given to the individual panels,(1) to (8), correspond to those at the nodes in the phylogenetic tree and are colored differentially to indicate comparable divergence times. Diagonal lines are colored according to the level of sequence similarity (dark green, 75-100%; light green, 50-74%; orange, 25-49%; yellow, 0-24%), and frequent repetitive sequences cause yellow dotted signals in the background. (*B*) Chromosomal homology suggested by synteny conservation of one-to-one ortholog pairs. See Methods for details. The color of the ribbons connecting the synteny represents each of the three categories of the zebra shark chromosomes: eMAC (gray); eMID (magenta); and eMIC (cyan). (*C*) Chromosomal homology between the zebra shark and other vertebrates. Conserved synteny is visualized with the inter-specific correspondence of one-to-one orthologs (see Methods).

Previous studies sought to reconstruct the process of karyotypic evolution of vertebrates but often lacked elasmobranchs in the dataset (Nakatani et al. 2021; Sacerdot et al. 2018). In the present study, we performed a gnathostome-wide comparison of the syntenic location of one-to-one orthologs, including the zebra shark (Figure 3B; Supplementary Figure 3). Remarkably, the majority of the genomic regions in the zebra shark eMIC were shown not to share one-to-one orthologs with the so-called microchromosomes of the chicken or spotted gar (Figure 3B). Moreover, the smallest spotted gar chromosomes were frequently shown to be homologous to zebra shark eMID (chromosomes 15–33), and not to its eMIC (chromosomes 34–) (Figure 3B). It was also shown that zebra shark chromosomes 10, 25, 26, 27, and 28 have chicken homologs of similar size but their *C. milii* homologs have been fused into larger chromosomes, while zebra shark chromosomes 2, 3, 4, 5, 6, 8, 11, 18, 20, 23, 33, 34, and 36 are likely to be products of fissions that occurred in the elasmobranch lineage (Figure 3C; Supplementary Figure 3). These results refute the common origins of microchromosomes among chicken, spotted gar, and zebra shark and indicate a more drastic reorganization of karyotypes at the base of jawed vertebrates than previously inferred from the comparison involving only the *C. milii* as a cartilaginous fish (Nakatani et al. 2021).

### Length-dependent properties of chromosomes

To characterize shark chromosomes in depth, base compositions, gene distributions, and molecular evolutionary rates quantified with genic synonymous substitutions (*K*_s_) were investigated (Figure 2B). As previously observed in birds and reptiles (Kuraku et al. 2006; Matsubara et al. 2012; Burt 2002; Waters et al. 2021; Srikulnath et al. 2021), relatively short zebra shark chromosomes tend to have higher GC-content, higher gene density, and smaller gene length than its larger chromosomes (Figure 2B). This chromosome length-dependent pattern is most pronounced in the chicken in which microchromosomes are canonically implicated (Figure 2B). Remarkably, this chromosome length-dependent pattern is observed not only in the spotted gar for which homology of microchromosomes to chicken was previously implicated (Braasch et al. 2016) but also in the western clawed frog *Xenopus tropicalis* which conventionally is thought to have no microchromosomes recognized cytogenetically (Uno et al. 2012) (Supplementary Figures 2).

Although this chromosome length-dependent pattern seems prevalent among phylogenetically diverse vertebrate species, some short chromosomes exhibit exceptionally low GC-content, such as chromosome 26 of the *C. milii*, chromosome 29 and 33 of the chicken, and Linkage Group 29 of the spotted gar (Figure 2; Supplementary Figures 1 and 2). These exceptions evoke a caution for the generalization of common characteristics of short chromosomes. It needs to be carefully examined whether the relatively short sequences with exceptionally low GC-content are fragments of large chromosomes that failed to be assembled to a chromosome scale.

### Do ‘macrochromosome’ ends resemble microchromosomes?

We analyzed regional variations within individual chromosomes of diverse vertebrates, including the zebra shark. To further characterize the distinct trend of chromosomal ends indicated in Figure 2, we separated the 1-Mb-long ends from relatively large chromosomes and analyzed the trends of genomic sequences as well as those of the coding genes (Figure 4A). Intact chromosome ends are known to be occupied by telomeric or subtelomeric simple repeats. Thus, we first examined whether the end of the chromosomal sequences used in this study harbor sufficient protein-coding genome segments that provide unambiguous trends. Our comparison detected a remarkable difference in GC-content of protein-coding regions between the 1-Mb-long chromosome ends and the remainder of the chromosome (Supplementary Figure 4), which resembled the pattern of the genomic sequences (Figure 4A). This observation suggests that the 1-Mb-long margins of the chromosomal DNA sequences employed in this study are not occupied solely by telomeric or subtelomeric simple repeats, but also engulf functional protein-coding genes that are affected by the trends of flanking genomic environments. This prompted us to analyze more genomic and genic properties of chromosome sequence ends. Our further comparison consistently revealed an increase of gene density and synonymous substitution rate, as well as a decrease in gene length, in the ends of relatively large chromosomes, compared with their remainders (Figure 4B). In both the zebra shark and chicken, the characteristics of chromosome ends resemble those of relatively small chromosomes (Figure 4B).

**Figure 4.**
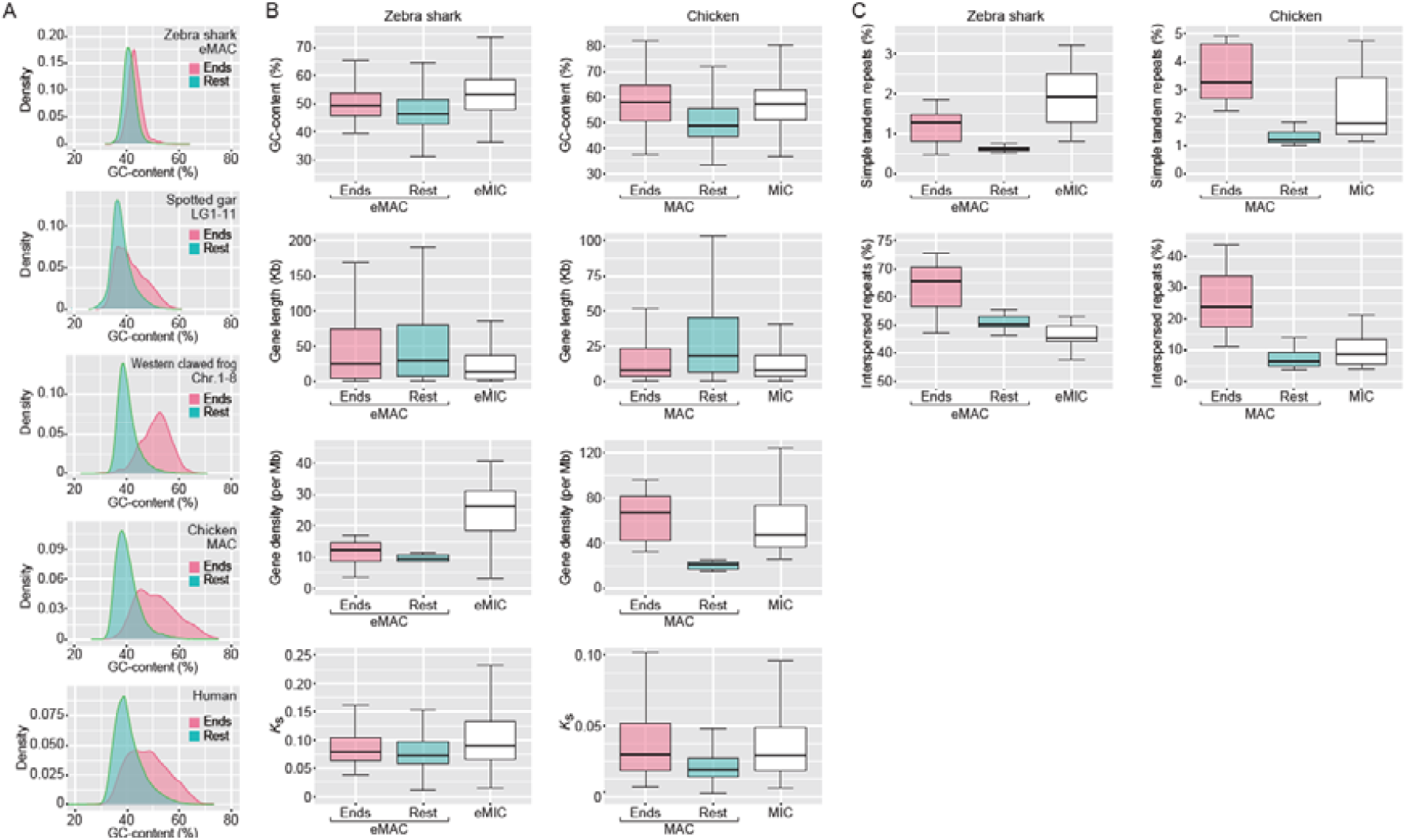
Intrachromosomal heterogeneity of sequence characteristics. (*A*) Comparison of global GC-content in 10 Kb-long non-overlapping windows between the 1-Mb-long ends and the remainders of relatively large chromosomes for diverse vertebrates. (*B*) Comparison of global GC-content of protein-coding regions, gene length, gene density, and synonymous substitution rate (*K*_s_) in 10 Kb-long windows among the 1 Mb-long chromosome ends, their remainders, and relatively small chromosomes (zebra shark eMIC and chicken microchromosomes (MIC)), for the zebra shark and chicken, respectively. (*C*) Differential distribution of simple tandem repeats and interspersed repeats. Proportions of the sequences identified as simple tandem repeats and interspersed repeats in 10 Kb-long windows were compared between the 1 Mb-long ends of relatively large chromosomes (zebra shark eMAC and chicken macrochromosomes (MAC)), their remainders, and relatively small chromosomes (zebra shark eMIC and chicken MIC).

### Intra-genomic repetitive element distribution

We also analyzed the distribution of repetitive elements on chromosomes of variable length. In fact, previous studies yielded equivocal observations. Some of those studies showed a higher abundance of repetitive elements on larger chromosomes (Koochekian et al. 2022), whereas others indicated localization biased toward smaller chromosomes (Hara et al. 2018). So far, no solid cross-species comparison has been made by taking the difference of repeat classes into account. In the newly obtained shark genome sequences, we separately quantified the sequence proportions identified as interspersed repeats (LINE, SINE, LTR, and DNA elements) and simple tandem repeats (simple repeats and low-complexity DNA sequences, including satellites). This resulted in contrasting patterns between these two repeat categories. Along with chromosome length increase, the interspersed repeat content increases, whereas the simple tandem repeat content decreases (Figure 2A; Supplementary Figure 1). Remarkably, the higher frequency of simple tandem repeats on smaller chromosomes is commonly observed in other vertebrates, including the chicken and the *C. milii* (Figure 2A).

As examined above for other characteristics, we dissected the observed chromosome length-dependent trend of repeat distribution, again by isolating the 1-Mb-long ends versus the remainder of relatively large chromosomes (Figure 4C). In this comparison, we observed a higher content of interspersed repeats in the ends of relatively large chromosomes, namely zebra shark eMAC and chicken MAC (Figure 4C). The higher repeat content was commonly observed in relatively small chromosomes (zebra shark eMIC and chicken MIC), except that interspersed repeat content is reduced in zebra shark eMIC (Figure 4C).

### First genomic characterization of a shark sex chromosome

So far, there has been no intensive DNA sequence-based characterization of sex chromosomes for chondrichthyan species. Expecting that a sex chromosome will exhibit a distinct male–female ratio of sequencing depth (Palmer et al. 2019), we performed short-read sequencing of the whole genomes for both sexes in zebra shark and whale shark. For these species, our previous cytogenetic analysis did not detect any heteromorphic sex chromosomes (Uno et al. 2020). Our comparison among different chromosomes detected a remarkably lower male-to-female sequencing depth ratio of close to 0.5 for chromosome 41 of both these species (Figure 5A). This suggests male as a heterogametic sex and the XY system for these species. Next, we investigated the origin of this putative X chromosome of these species by locating human and chicken orthologs of the genes on shark chromosomes (Figure 5B, magenta). This revealed chromosome-level homology of the putative X chromosomes of the zebra shark to a part of human chromosome 12 and chicken chromosome 34 (Figure 5B, magenta). Neither the human X, human Y, chicken Z, nor chicken W chromosomes exhibited pronounced homology to the putative zebra shark X chromosome. The putative shark X chromosome identified in this study do harbor orthologs of a number of well-studied regulatory factors but do not harbor the orthologs of the sex determination genes identified in other vertebrates, Dmrt1- or Sox3-related transcription factors as well as components of TGF-β signaling pathway, such as Amh, Amhr2, Bmpr1b, Gsdf, and Gdf6 (Bertho et al. 2021).

**Figure 5.**
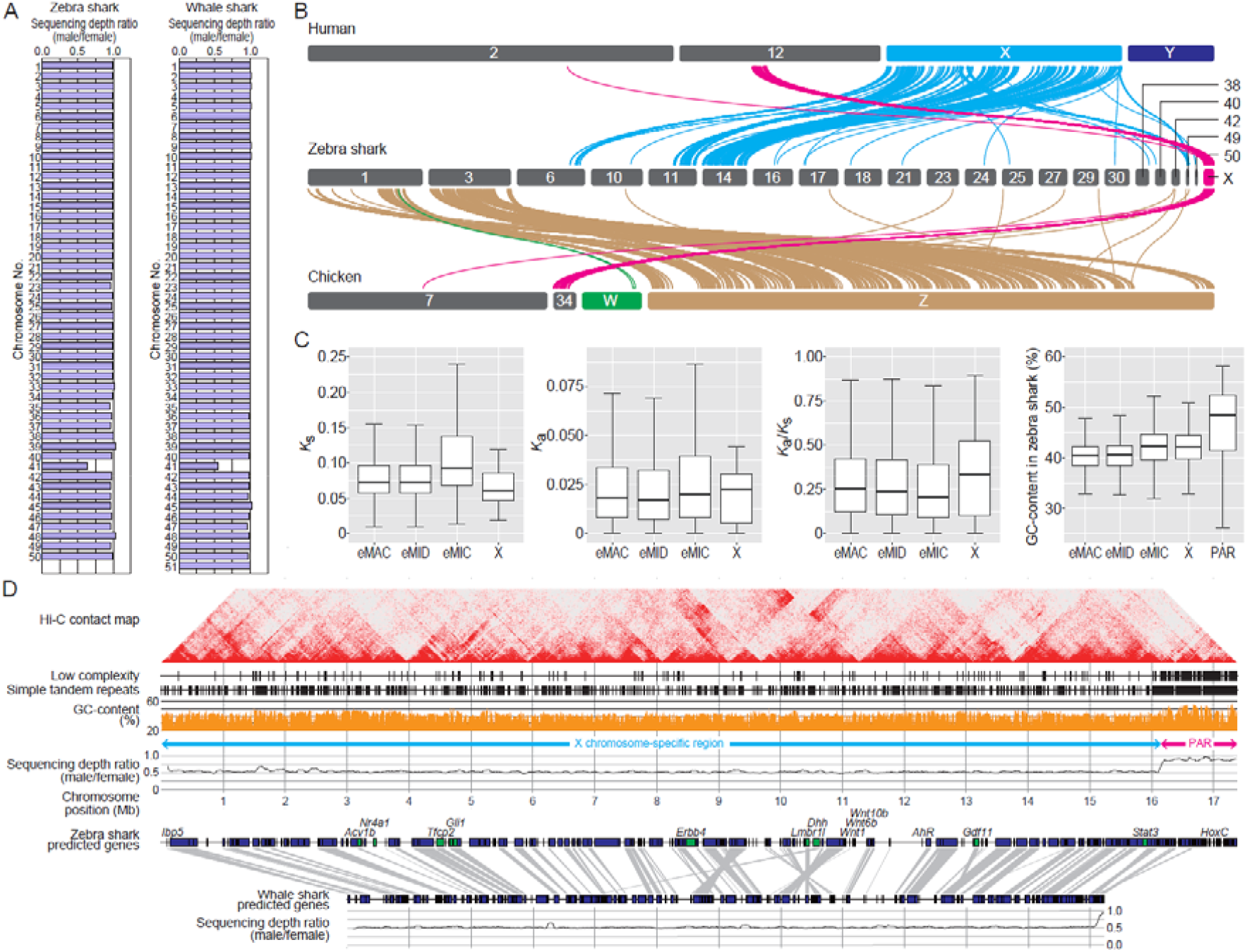
Genomic identification of the zebra shark X chromosome. (*A*) Male–female ratio of short-read sequencing depth in the shark chromosomes. (*B*) Cross-species synteny of sex chromosome-linked genes based on 1-to-1 orthologs. (*C*) Characteristic comparisons between the different chromosome categories in the zebra shark. *K*_s_ (synonymous substitution rate) and *K*_a_ (nonsynonymous substitution rate) values were calculated for 1-to-1 ortholog shared with the whale shark. Only for GC-content, the PAR was shown separately, because it contained few genes. (*D*) Structural comparison between the zebra shark and whale shark X chromosome sequences.

In the approximately 17 Mb-long sequence of the putative zebra shark X chromosome, one 1.5 Mb-long end exhibited a male–female sequencing depth ratio of nearly 1.0 (Figure 5D). This region, with a sequence depth comparable to that of the autosomes, is deduced to be a pseudoautosomal region (PAR) that is likely shared between the heterogametic sex chromosomes X and Y (Smeds et al. 2014; Palmer et al. 2019). We characterized possible unique patterns of molecular evolution typical of sex chromosomes (Figure 5C). In the X chromosome, protein-coding genes exhibited a relatively low synonymous substitution rate (*K*_s_) and high non-synonymous substitution rate (*K*_a_), resulting in an increased *K*_a_/*K*_s_ ratio. Our comparison also revealed higher frequency of low-complexity repeats as well as higher GC-content in the PAR (Figure 5C and D).

Our comparison of the ortholog location between the putative X chromosomes of the two species supported a high cross-species conservation of the chromosome structure (Figure 5D). In the current whale shark assembly, chromosome 41, which corresponds to the putative X chromosome, may not cover the whole chromosome, possibly excluding one end of the PAR (Figure 5D).

### Identification of Hox C cluster on the putative X chromosome

In the newly obtained sequence of the putative zebra shark X chromosome, we identified an array of orthologs of the non-shark genes encoding homeobox proteins Hox C (Figure 5D). In elasmobranchs, Hox C genes were long thought missing from the genome (King et al. 2011) but later identified in several shark species, as rogue open reading frames (ORFs) flanked by massively repetitive sequences (Hara et al. 2018; Kuraku 2021). Our genome-wide gene prediction for the zebra shark detected the ORF of *Hoxc8, -c11*, and *-c12*, while the partial ORF of the putative *Hoxc6* ortholog was also identified by a manual search of the raw genomic sequence (Figure 6A). These Hox C genes were located in a 180 Kb-long genomic segment in the PAR of the putative X chromosome (Figure 6A), identified as a single cluster in an elasmobranch fish for the first time. Their orthologies were confirmed with molecular phylogenetic trees, which also indicated elevated molecular evolutionary rates with long branches for elasmobranch Hox C genes (Figure 6B). Our RNA-seq data showed the transcription of these Hox C genes (except for *Hoxc12*) in embryos and juvenile tissues (Figure 6C). The identified zebra shark Hox C cluster is massively invaded by repetitive elements unlike the other Hox gene clusters (A, B, and D clusters) of this and many other vertebrate species (Figure 6A), as in the Hox C-containing genomic segments of other shark species (Hara et al. 2018). Our search for zebra shark orthologs of the protein-coding genes located near the human Hox C cluster (e.g., ATF7, CBX5) revealed poor conservation of the gene compositions. Some of the zebra shark orthologs were not identified in its entire genome sequence, suggesting a divergent nature of the genomic regions flanking the Hox C cluster.

**Figure 6.**
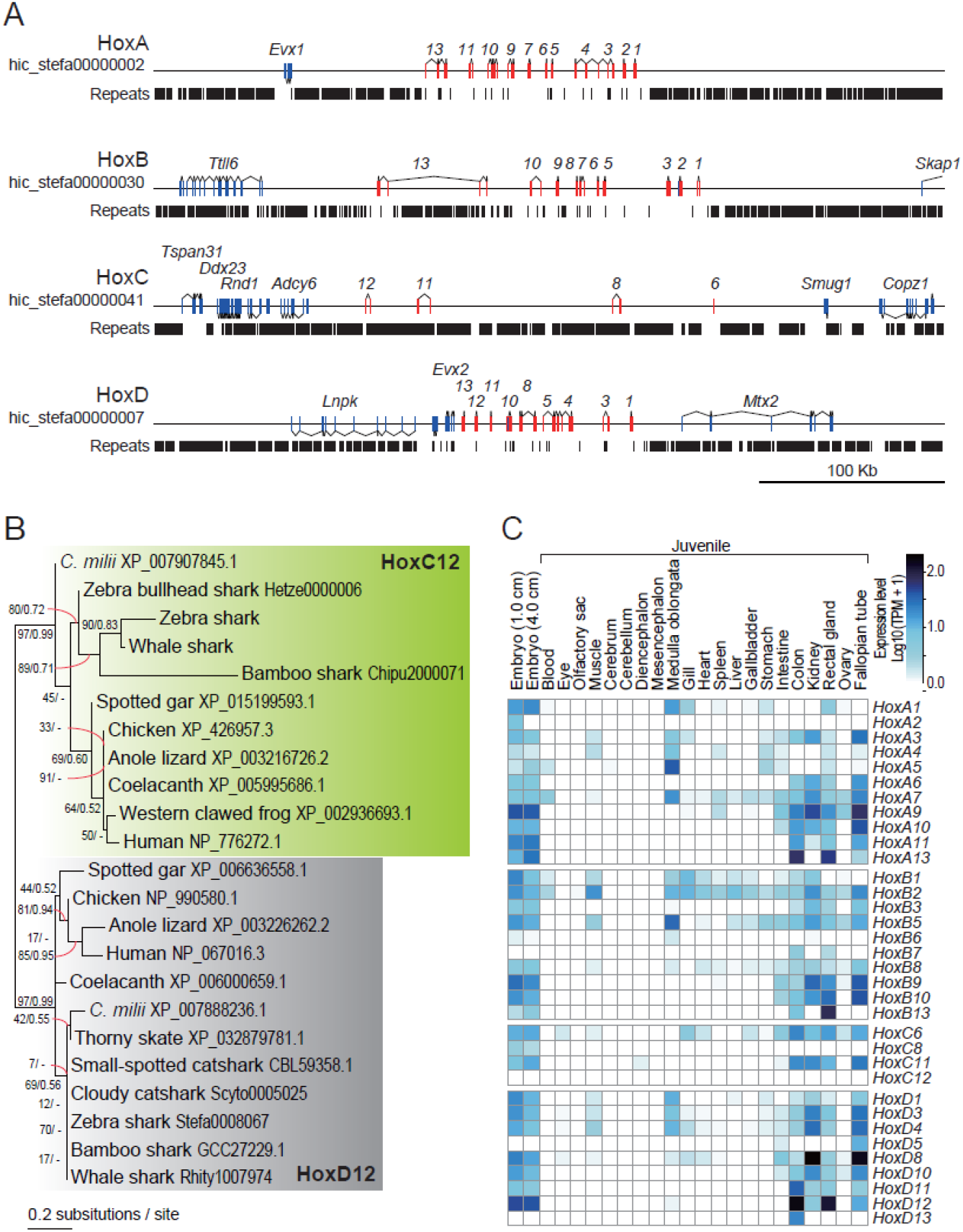
Zebra shark Hox C genes. (*A*) Genomic structure of the zebra shark Hox clusters and their neighboring regions. The exons of the Hox genes are shown in red boxes. (*B*) Molecular phylogenetic tree of Hox12 group of genes. The tree was inferred with the maximum-likelihood method as described in Methods. (*C*) Expression profiles of the zebra shark Hox genes in embryos and various tissues of a juvenile.

## Discussion

In this study, we chose two orectolobiform shark species (zebra shark and whale shark) in Elasmobranchii and characterized their genomic organization with chromosome-scale DNA sequences. This study was achieved in support of epigenome and transcriptome data prepared using fresh tissue samples and previously obtained karyotype information. Our results suggest that their karyotypes are organized by chromosomes of gradual sizes marked with size-dependent sequence properties (Figure 2). The pattern in the shark chromosomal organization is unique in its diversity among vertebrates (Uno et al. 2020), which is characterized by abundant chromosomes (up to 106 for diploids) and variable chromosome sizes. The abundance and highly variable sizes of chromosomes are known for some avian species, but the shark genome organization is distinct from avian counterparts in that shark chromosomes generally have higher repeat content than those of the chicken (Figure 2). In our comparison, the shark karyotype is also characterized by the ratio of the largest and smallest chromosome lengths of 40 to 100, compared with < 10 for most vertebrates, except for species with microchromosomes (Figure 1A). In fact, the length of the shortest chromosomal sequence in typical genome assemblies deposited currently in public databases is often dependent on sequence length cutoff by researchers’ decision. Especially for species with no solid karyotypic reference such as *Callorhinchus milii*, the range of sequences considered as chromosomes needs to be carefully examined.

Our chromosome-scale genome sequencing and analysis were enabled by access to fresh tissue samples. Because of low accessibility, no previous efforts could provide a set of DNA sequences, karyotypic configuration, and reliable measure of nuclear DNA content for a single chondrichthyan species. Of these, the two latter elements serve as indispensable references to validate the output of sequencing. These requirements are satisfied for both zebra shark and whale shark in our study. Especially for the whale shark, no published studies employed chromatin contact data for Hi-C scaffolding and transcriptome sequencing (Tan et al. 2021; Marra et al. 2019; Weber et al. 2020). In our study, the access to embryos and blood not only enabled chromosome-scale genome scaffolding but also provided transcriptional evidence of most shark Hox C genes that have been shown to exist in a cluster for the first time (Figure 6).

Peculiar fractions of vertebrate karyotypes with small sizes and higher GC-content have traditionally been designated as microchromosomes, which usually denote chromosomes shorter than 20 Mb (e.g., Nakatani et al. 2021) but have no uniform definition (see Introduction). Our investigation, focusing on various aspects of chromosomal DNA sequences, provided a novel view of vertebrate karyotypes that cannot be understood with a simple binary classification, namely with or without microchromosomes, or between macro- and microchromosomes. This view is supported by the common pattern of high intrachromosomal heterogeneity within individual macrochromosomes (Figure 4; Supplementary Figure 4) as well as interchromosomal heterogeneity among different microchromosomes (Figure 2; Supplementary Figures 1 and 2). In particular, the heterogeneity within macrochromosomes, marked with high GC-content, high gene density, and small gene length of their ends, may be shared widely among diverse vertebrates (Figure 4A, B). Intragenomic heterogeneity of GC-content was previously suggested to be caused by GC-biased gene conversion (Mugal et al. 2015). In addition, the uniform numbers of recombination per chromosome (Dumont and Payseur 2008) have been thought to explain the chromosome length-dependent GC-content variation between different chromosomes. Our observations did not show pronounced length-dependent variation of GC-content for chromosomes that were longer than 100 Mb (Figure 2B; Supplementary Figure 2). Taken together, we speculate that the peculiar nature of chromosome ends mainly account for the variation of GC-content among different chromosomes (Figure 4B). Importantly, ‘chromosome ends’ in this context not only harbor telomeric or other simple repeats but also hold complex sequences including protein-coding genes in sequence stretches that are longer than 1 Mb (Figure 4B; Supplementary Figure 4). In the *X. tropicalis* genome, such sequence stretches marked with elevated GC-content span much longer ranges than in other genomes (Supplementary Figure 1 and 2). The observed features of smaller chromosomes with larger proportions of such ‘ends’ in length are more affected than those of longer chromosomes, which likely explains the length-dependent nature of chromosomes. The nature of macrochromosome ends (e.g., with higher GC - content) is thought to be a remnant of the fusion of one or more microchromosome(s) to a macrochromosome (Waters et al. 2021). This hypothesis is not supported by our observation that even species possessing no explicit microchromosomes (e.g., *Xenopus tropicalis*) have chromosome ends with the peculiar nature of DNA sequences (SupplementaySupplementary Figures 1, 4). Remarkably no close relatives of *X. tropicalis* have been shown to possess microchromosomes),(Tymowska 1991), and thus microchromosome fusions cannot account for the characteristics of their chromosome ends with higher GC-content.

Our genome-wide sequencing depth investigation covering both sexes revealed the XY system for the two studied shark species and enabled the first sequence-based identification of shark sex chromosomes (namely, X chromosomes; Figure 5A). The genes on the shark X chromosome tend to exhibit larger non-synonymous substitution rates (*K*_a_) and smaller synonymous substitution rates (*K*_s_) than those on autosomes, resulting in a higher *K*_a_/*K*_s_ ratio (Figure 5D). This resembles the pattern known in other species including birds and insects, which is known as the ‘Faster X (Faster Z) hypothesis’ (Meisel and Connallon 2013; Charlesworth et al. 2018; Xu et al. 2019; Mank et al. 2007). The smaller *K*_s_ value is also indicative of the support for the male-driven hypothesis (Li 2002; Miyata et al. 1987), which is also to be examined by higher *K*_s_ values for genes on the presumptive Y chromosome even though it remains unidentified. Our comparison of protein-coding gene compositions showed that the zebra shark and the whale shark largely share the X chromosome that is homologous to each other (Figure 5D). The X chromosome harbors the Hox C cluster that was previously shown to be highly divergent and degenerative, but its localization in the PAR (Figure 5D) suggests a balanced dosage between males and females.

The whale shark is known as the largest extant ‘fish’, and one of its extant closest relatives is the zebra shark (Naylor et al. 2012). In elasmobranch evolution, the lineages leading to these two species diverged no earlier than 48.6 million years ago (Long 1992). The remarkably high similarity of the chromosome-scale sequence organization (panel 1 in Figure 3; Supplementary Figure 3) as well as the gene compositions on the X chromosome between these two species (Figure 5D) indicates a lower rate of chromosomal rearrangement in these lineages compared with those of species pairs of similar divergence times in other vertebrate lineages (Figure 3A). Among vertebrates, mammals and birds have relatively long-standing sex chromosomes (X/Y and Z/W, respectively) shared throughout these individual taxa that arose more than 48.6 million years ago (Long 1992). Although still limited in number, sex chromosomes have been identified in some elasmobranch species by cytogenetic analyses, all of which have a male-heterogametic system (Uno et al. 2020), as shown for the zebra shark and whale shark with genome sequencing in this study. Sharks, or a phylogenetically wider subset of cartilaginous fishes, may possess even older sex chromosomes, depending on phylogenetic prevalence of their homologs in more distantly related shark and even ray species.

Our synteny analysis yielded novel insights into genome evolution encompassing the whole diversity of vertebrates. It showed homology of the shark X chromosome to human chromosome 12 and chicken chromosome 34 (Figure 5B), which suggests an independent origin of the sex chromosome in elasmobranchs. The synteny analysis involving sharks also provided clues to the origin of microchromosomes. It suggests that eMIC, the shark’s small chromosomes, are homologous to several of the large chromosomes in both chicken and spotted gar and are not necessarily homologous to their microchromosomes (Figure 3B). Also, microchromosomes of the chicken and gar were not shown to be homologous to eMICs (Figure 3B). It is crucial for any effort to reconstruct the diversity of chromosome organization in vertebrates to incorporate elasmobranchs into the comparisons.

## Methods

### Animals

Fresh blood from a female adult zebra shark (total length, 2.2 m); Individual ID, sSteFas1 and a male adult whale shark (total length, 8.8 m; Individual ID, sRhiTyp1) were sampled at Okinawa Churaumi Aquarium and used for the preparation of whole-genome shotgun DNA libraries, Hi-C libraries, and RNA-seq libraries as well as for measuring nuclear DNA content by flow cytometry. Likewise, fresh blood of a male zebra shark (total length, 2.1 m; Individual ID, sSteFas2) and a female whale shark (total length, 8.0 m; Individual ID, sRhiTyp2) were sampled and used for quantifying the male/female ratios of individual chromosomal regions. Extraction of ultra-high molecular weight DNA was performed by collecting blood cells by centrifugation, and the collected cells were embedded in agarose plugs (4.0 × 10^5^ cells/plug). The agarose gel plugs were prepared and processed with the CHEF Mammalian Genomic DNA Plug Kit (BioRad, Cat. No. #1703591). Total RNAs used to construct RNA-seq libraries were extracted from various tissues of a female juvenile zebra shark (total length, 30 cm) born at Okinawa Churaumi Aquarium and a female juvenile whale shark (total length, 7.7 m; Individual ID, sRhiTyp3) (Supplementary Table 3). These animals were introduced into the aquarium in accordance with local regulations before those species were assessed as endangered. Animal handling and sample collections at the aquarium were conducted by veterinary staff without restraining the individuals under the experiment ID AT19002 approved by Institutional Animal Care and Use Committee of Okinawa Churashima Foundation in accordance with the Husbandry Guidelines approved by the Ethics and Welfare Committee of Japanese Association of Zoos and Aquariums. All other experiments were conducted in accordance with the Guideline of the Institutional Animal Care and Use Committee (IACUC) of RIKEN Kobe Branch (Approval ID: H16-11).

### Genome sequencing and scaffolding

For a female zebra shark, paired-end and mate-pair DNA libraries for *de novo* genome sequencing were prepared and sequenced as previously described (Hara et al. 2018; Yamaguchi et al. 2021). The amount of starting DNA and numbers of PCR cycles for the library preparation are included in Supplementary Table 3. The total sequencing coverage amounted to 95.8 times the genome size based on the reference measured previously by flow cytometry (3.71 Gb; Kadota et al., in preparation). Low-quality bases from paired-end reads were removed by TrimGalore v0.6.6 with the options ‘--stringency 2 --quality 20 --length 25 --paired -- retain_unpaired’. As described previously (Hara et al. 2018), short-read assembly of the zebra shark, as well as scaffolding with mate-pair reads followed by gap closure, was performed using Platanus v1.2.4 (Kajitani et al. 2014).

Whole genome sequencing for a male whale shark employed the 10x Genomics Chromium to produce Linked-Read data. A DNA library was prepared using 12 ng of gDNA extracted from blood cells according to the user guide of the Chromium Genome Library Kit v2 Chemistry using the Chromium Genome Library Kit & Gel Bead Kit v2 (10x Genomics, Cat. No. #120258) and the Chromium Genome Chip Kit v2 (10x Genomics, Cat. No. #120257). The library was sequenced on a HiSeq X (Illumina) platform to obtain 151 nt-long paired-end reads. Sequence assembly using the Linked-Read data of 46.4 times the genome size was performed with the program Supernova v2.0. The resultant sequences were subjected to scaffolding with the program P_RNA_scaffolder (commit 7941e0f in GitHub) (Zhu et al. 2018) using the result of the alignment of transcriptome sequence reads (obtained as described below) performed with the program HISAT2 v2.1.0 onto those genome sequences (Kim et al. 2019).

### Hi-C data production and chromosome-scale genome scaffolding

Hi-C libraries of the zebra shark and whale shark were constructed using restriction enzymes DpnII and HindIII, respectively, as previously reported (Kadota et al. 2020). Blood cells collected as described above were fixed in 1% formaldehyde solution. A fixed tissue containing 10 μg of DNA was used for the preparation of Hi-C DNA via *in situ* restriction digestion and ligation. The Hi-C library was prepared using 2 μg of the ligated DNA with five cycles of PCR amplification. Quality controls of the ligated DNA and the Hi-C libraries were performed as described previously (Kadota et al. 2020).

Each of the zebra shark and the whale shark genome assemblies was used for Hi-C read mapping with Juicer v1.5 and chromosome-scale scaffolding with the program 3d-dna (v180922) (Dudchenko et al. 2017). In the scaffolding, three different lengths were tested (5, 10, and 15 kb) for the option ‘-i’ defining the input sequence length threshold. For each species, the three resulting scaffolding outputs, as well as the original assembly before Hi-C scaffolding, were assessed based on sequence length distribution and protein-coding gene completeness. Among all the scaffolding outputs compared, the output with the option ‘-i 10000’ was judged to be optimal and was subjected to a “review” of the scaffolding results on Juicebox v1.11.08 (Durand et al. 2016) to minimize inconsistent signals of chromatin contacts (Supplementary Figure 5). The review was facilitated by referring to nucleotide sequence-level similarity between different scaffolding outputs visualized by SyMAP v5.0 (Soderlund et al. 2011). After the review, the sequences judged as contaminants from other organisms were removed, as previously reported (Hara et al. 2018). The resulting sequences were deposited as JAHMAH000000000 and JAFIRC000000000 in DDBJ/NCBI under BioProject PRJNA703743.

### Repeat identification

To obtain a species-specific repeat library, RepeatModeler v2.0.2a was run on the genome assembly of the individual species with default parameters (Smit and Hubley 2008). Detection of repeat elements in the genome was performed by RepeatMasker v4.1.2-p1 (Smit et al. 2013) with RMBlast v2.6.0+, using the species-specific repeat library obtained above. For quantification of the content of interspersed repeats and simple tandem repeats, RepeatMasker was run separately with the options ‘-nolow -norna’ and ‘-noint -norna’, respectively.

### Gene model construction

The program Braker v2.1.6 was employed for gene prediction by inputting the results of RNA-seq read mapping to a genome assembly in which repetitive sequences are soft-masked by RepeatMasker with the options ‘-nolow -xsmall’, as well as amino acid sequences of closely related species as homolog hints (Brůna et al. 2021; Smit et al. 2013). To build the homolog hints based on the amino acid sequences, we used the previously reported amino acid sequence sets of the brownbanded bamboo shark and the whale shark.

Gene space completeness in Figure 1C was obtained by the BUSCO pipeline ver. 5 (Seppey et al. 2019) using the BUSCO’s Vertebrata ortholog set.

### Synonymous and non-synonymous substitution quantification

In order to calculate the number of synonymous substitutions per synonymous site (*K*_s_) and the number of nonsynonymous substitutions per non-synonymous site (*K*_a_), the 1-to-1 orthologs shared by the four elasmobranch species (zebra shark, whale shark, brownbanded bamboo shark, and cloudy catshark) were selected by SonicParanoid v1.3.4 (Cosentino and Iwasaki 2019) as follows. First, peptide sequences of the retrieved orthologs were aligned with MAFFT v7.475 with the option ‘-linsi’ (Katoh and Standley 2013). The individual alignments were trimmed and back-translated into nucleotides with trimal v1.4.rev15 with the options ‘-automated1 -backtrans’ followed by the removal of gapped sites using trimAl with the option ‘-nogaps’ (Capella-Gutiérrez et al. 2009). Ortholog groups containing fewer than 100 aligned codons or a stop codon were discarded. For the selected ortholog groups, *K*_s_ and *K*_a_ were computed with yn00 in the PAML v4.9c88 (Yang 2007). Computed values larger than 0.01 and smaller than 99 were included in the results (Figures 2B, 4B, and 5C).

### RNA-seq and transcriptome data processing

Total RNAs were extracted with Trizol reagent (Thermo Fisher Scientific). Quality control of the RNA treated with DNase I was performed with Bioanalyzer 2100 (Agilent Technologies). Libraries were prepared with TruSeq RNA Sample Prep Kit (Illumina) or TruSeq Stranded mRNA LT Sample Prep Kit (Illumina) as previously described (Hara et al. 2018). The amount of starting total RNA and numbers of PCR cycles are included in Supplementary Table 3. To remove adaptor sequences and low-quality bases, the obtained sequence reads were trimmed with TrimGalore v0.6.6 as outlined above, and *de novo* transcriptome assembly was performed with the program Trinity v2.11.0 with the option ‘--SS_lib_type RF’ (Grabherr et al. 2011). The trimmed RNA-seq reads were aligned to the genome assembly using the program HISAT2 v2.1.1 (Kim et al. 2019), which was followed by gene expression quantification with StringTie v2.0.6 (Pertea et al. 2016).

### Conserved synteny detection

Characterization of chromosomal homology among different species employed predicted protein sequence datasets available at the NCBI RefSeq database. After alternative splicing variants were removed, one-to-one orthologs were selected by SonicParanoid v1.3.4 with the option ‘most-sensitive’ (Cosentino and Iwasaki 2019). Conserved synteny between species was visualized based on single copy 1-to-1 orthologs using RIdeogram (Hao et al. 2020).

### Molecular phylogeny inference

Amino acid sequences were retrieved from aLeaves (Kuraku et al. 2013). Multiple sequence alignment was performed with MAFFT with the option ‘-linsi’. The aligned sequence sets were processed using trimAl v1.4 rev15 with the option ‘-automated1’ (Capella-Gutiérrez et al. 2009). This was followed by another trimAl run with the option ‘-nogaps’. Molecular phylogenetic trees were inferred by RAxML with the ‘-m PROTCATWAG -f a -# 1000’ options unless stated otherwise (Stamatakis 2014). Tree inference in the Bayesian framework was performed with the program PhyloBayes v4.1c with the options ‘-cat -dgam 4 -wag -nchain 2 1000 0.3 50’ unless stated otherwise. This was followed by an execution of bpcomp in the PhyloBayes v4.1c package with the option ‘-x 100’ (Lartillot et al. 2009). The support values at the nodes of molecular phylogenetic trees included are, in order, bootstrap values and Bayesian posterior probabilities. The latter was shown only when the relationship at the node in the visualized tree was supported by the Bayesian inference.

### Identification of the X chromosome

To identify a chromosome-scale scaffold with a distinct male–female sequencing depth ratio in the zebra shark, the same number of trimmed genomic shotgun reads (293,686,584) was prepared for both sexes. The reads were mapped with BWA-mem2 (v2.2.1) onto a genome assembly (Li and Durbin 2009). Mapped reads were counted for male and female using the bamtobed program in the package bedtools v2.29.2, for each scaffold in 10 Kb non-overlapping windows, and male–female ratios were calculated (Quinlan and Hall 2010). Sequencing depth of male and female reads on the identified X chromosome was also calculated for 10 Kb non-overlapping windows. Windows with the proportion of ambiguous bases of more than 50% were excluded from computation. This procedure was also applied to the whale shark, using 220,785,436 trimmed reads.

## Data access

The sequence data resulting from the present study were deposited as JAHMAH000000000 and JAFIRC000000000 in DDBJ/NCBI under BioProject PRJNA703743.

## Acknowledgments

We thank Tomoyuki Furuyashiki and Masayuki Taniguchi for their cooperation in obtaining Linked-Read sequence data, Chiharu Tanegashima and Kaori Tatsumi for their support in sequence data acquisition, animal caretakers including Yano Nagisa at Okinawa Churaumi Aquarium for their assistance, and Yukiko Imai for valuable discussion. Computations were partially performed on the NIG supercomputer at ROIS National Institute of Genetics. This study was funded by intramural budgets granted by RIKEN and the National Institute of Genetics, as well as JSPS KAKENHI Grant Nos. 20H03269 and 22K15088.

